# Structure and engineering of miniature *Acidibacillus sulfuroxidans* Cas12f1

**DOI:** 10.1101/2023.07.03.547593

**Authors:** Zhaowei Wu, Dongliang Liu, Deng Pan, Haopeng Yu, Jin Shi, Jiacheng Ma, Wenhan Fu, Zhipeng Wang, Zijie Zheng, Yannan Qu, Fan Li, Weizhong Chen, Xingxu Huang, Huaizong Shen, Quanjiang Ji

## Abstract

The miniature CRISPR-Cas12f nucleases allow for efficient delivery via cargo-size-limited adeno-associated virus delivery vehicles, thereby showing promising potential for in vivo therapeutic applications. *Acidibacillus sulfuroxidans* Cas12f1 (AsCas12f1, 422 amino acids) is the most compact Cas12f nuclease identified to date, showing a moderate level of genome-editing activity in human cells compared to Cas9 and Cas12a. Understanding the mechanisms of why such a compact nuclease is active for genome editing would facilitate its rational engineering. Here, we report the cryo-EM structure of the AsCas12f1-sgRNA-dsDNA ternary complex, and reveal that AsCas12f1 functions as an asymmetric dimer for single-guide RNA (sgRNA) binding and DNA targeting. The detailed mechanisms for dimer formation, PAM recognition, and sgRNA accommodation are elucidated. Leading by the structural knowledge, we extensively engineer the AsCas12f1 nuclease and its corresponding sgRNA, resulting in an evolved AsCas12f1-sgRNA combination with drastically enhanced genome editing activity in human cells. These results provide further understanding of compact CRISPR systems and expand the mini CRISPR toolbox for therapeutic applications.

## Introduction

The clustered regularly interspaced short palindromic repeats (CRISPR) and CRISPR-associated (Cas) system provide prokaryotes and some huge bacteriophages adaptive immunity against invasive nucleic acids^1^. CRISPR-Cas systems are diverse in their genetic compositions and functional configurations, which are currently divided into two distinct classes (class 1 and 2) and six major types (type I-VI)^1, 2^. The class 2 systems, consisting of type II, V, and VI, are distinguished by the RNA-guided single multi-domain effector nucleases^1, 2^, such as Cas9, Cas12, and Cas13, and have been engineered into groundbreaking genome-editing tools^3, 4^. Among them, Cas9 and some Cas12 nucleases induce double-stranded breaks (DSBs) to the genomic DNA, resulting in insertion and deletion (indel) mutations via the nonhomologous end-joining (NHEJ) mechanism^5^, or precise genetic modifications using homology-direct repair (HDR) with the supplementation of repair templates^5^. In addition, the fusion with deaminases creates base editors that change DNA in a single-base-pair precision^6–8^, and the combination with reverse transcriptase generates prime editors, allowing the introduction of any pre-determined type of short insertion, deletion, and mutation^9^. These technologies substantially facilitate the genetic engineering in various organisms^5, 10, 11^, and offer new therapeutic options to treat genetic disorders^12^, cancers^13^, and infectious diseases^14^. However, the most commonly-used Cas9 and Cas12a nucleases are large in their molecular sizes (mostly greater than 1,000 amino acids), posing difficulties in efficient cellular delivery, particularly in scenarios that uses cargo-size-limited delivery vehicles, such as adeno-associated virus (AAV)^15^.

Recent studies identified a set of compact CRISPR nucleases that are capable of dsDNA targeting, including Cas12e (also known as CasX, ∼1,000 a.a.)^16^, Cas12f (formerly Cas14, 400-700 a.a.)^17–20^, Cas12i (∼1,000 a.a.)^21–24^, Cas12j (or CasΦ, 700-800 a.a.)^25–27^, Cas12l (or Casπ, ∼860 a.a.)^28^, Casλ (∼800 a.a.)^29^, TnpB (∼400 a.a.)^30^, and IscB (∼400 a.a.)^31, 32^. However, most of them are either incompetent or inefficient for genome editing in mammalian cells with their natural forms^16, 25, 30, 31, 33–35^. Through structure-guided engineering, some of them are transformed into robust and efficient genome editors^36–40^. Among these compact CRISPR nucleases, Cas12f appears to strike the perfect balance between genome editing efficient and molecular size at this moment. The first identified Cas12f nuclease, Un1Cas12f1 (also known as Cas14a1, 529 a.a.) from uncultivated archaeon, was initially found to target ssDNA exclusively^41^. Later, it was confirmed to be capable of dsDNA cleavage^17, 18^. However, the natural form of Un1Cas12f1 merely showed an indel activity at a negligible level on limited endogenous loci in eukaryotic cells^33^.

The cryo-EM structure study of the Un1Cas12f1-sgRNA-dsDNA complex revealed an asymmetric homodimer configuration for binding sgRNA to target a dsDNA substrate^19, 20^. Intriguingly, nearly half of the natural gRNA scaffold is flexible in the structure^19, 20^. Through systemically truncating the flexible region in sgRNA, Un1Cas12f1 was transformed into an efficient and specific genome editor in mammalian cells^36, 37^. In contrast, our previous study showed that *Acidibacillus sulfuroxidans* Cas12f1 (AsCas12f1, 422 a.a.) is a naturally active genome editor^17^, presenting a moderate level of indel activity relative to the widely used *Streptococcus pyogenes* Cas9 (SpCas9) and *Acidaminococcus sp.* Cas12a (AsCas12a). However, similar truncations in the sgRNA counterparts of AsCas12f1 either partially or completely abolished its dsDNA cleavage activity^17^. These findings imply that despite the sequence similarity (∼26% identity, and ∼45% positive) between the two Cas12f1 nucleases, AsCas12f1 may distinguish from Un1Cas12f1 by its structural configurations, making it active as a genome editor with an even smaller molecular size.

In this article, we report the cryo-EM structure of the AsCas12f1-sgRNA-dsDNA ternary complex at 2.7-Å resolution, revealing that asymmetrically dimerized AsCas12f1 proteins accommodate the sgRNA for dsDNA targeting. Through comparative structural analysis against Un1Cas12f1, the structure uncovers several unique structural features within the dimer interface, the PAM recognition region, and the AsCas12f1 sgRNA, together explaining the key mechanism of how AsCas12f1 is active as a genome editor with such a small molecular size. Next, the AsCas12f1 nuclease and its corresponding sgRNA are extensively optimized by structure-guided engineering, finally resulting in an evolved genome editor (AsCas12f1-v5.1 + sgRNA_T1) with substantially enhanced genome-editing activity. These findings lay the groundwork for understanding the fundamental working mechanism and the molecular evolution of mini-size CRISPR nucleases, providing practical knowledge for improving the performance of genome editors, and expanding the CRISPR genome-editing toolbox for therapeutic applications.

## Results

### AsCas12f1 dimer binds a sgRNA for DNA targeting

To uncover the underlying mechanisms for AsCas12f1-mediated dsDNA cleavage, we determined the cryo-EM structure for the ternary complex consisting of the nuclease-deactivated AsCas12f1 variant (D225A) with its cognate 191-nt sgRNA and the target dsDNA (30-nt TS + 6-nt NTS) with a 5’ GTTG PAM, at an overall resolution of 2.7 Å (Fig. 1a-e, Supplementary Fig. S1a-e, and Table S1). The high-resolution map allowed us to build a detailed atomic model for the ternary complex (Supplementary Fig. S2). As shown in Fig. 1c-e, the C-shaped whole nuclease consists of two AsCas12f1 molecules, designated as AsCas12f1.A and AsCas12f1.B that form an asymmetric homodimer, and bind to a sgRNA molecule for target dsDNA recognition. An AsCas12f1 monomer can be divided into an amino-terminal domain (NTD) and a carboxy-terminal domain (CTD) which are connected by a linker loop (Fig. 1a). The NTD comprises a recognition domain (REC) and a wedge domain (WED). The CTD consists of a RuvC domain and a zinc-finger domain (ZF). The two monomers of AsCas12f1 interact extensively with each other through their REC domains (Fig. 1c-e). The map density of the complete B.CTD in AsCas12f1.B is missing, indicating its structural flexibility (Fig. 1a, c-e). Despite its small molecular size, the dimeric AsCas12f1 nuclease resembles a distinct miniature bilobed structure consisting of a recognition (REC) lobe and a nuclease (NUC) lobe (Fig. 1c). The REC lobe is formed by the A.REC, A.WED, B.REC, and B.WED domains. The NUC lobe is comprised by the A.RuvC and A.ZF domains. The whole nuclease clamps the heteroduplex of the target DNA and the guide sequence through the central groove between the REC and NUC lobes (Fig. 1c-e). The PAM duplex is recognized by A.REC and A.WED. The AsCas12f1-cognate sgRNA consists of a 171-nt sgRNA scaffold and a 20-nt guide sequence, forming six stems (stem 1∼4, and two repeat:anti-repeat segments) with a sophisticated tertiary RNA structure (Fig. 1b-e). Interestingly, the sgRNA scaffold is stacked with the whole nuclease side by side (Fig. 1c-e). The stem 2, 4, and R:AR-2 predominantly interact with AsCas12f1, whereas the stem 1, 3, and R:AR-1 barely does. Phylogenetic analysis and structural comparison of AsCas12f1 with Un1Cas12f1 reveal major structural differences in dimer configuration and sgRNA architecture, despite of their close sequence similarity (Supplementary Fig. 3a-c). The ISDra2 TnpB is considered as an ancestor of most Cas12 nucleases^30, 42, 43^. Recent structural studies revealed that TnpB shares similar domain configurations as Cas12f nucleases, but TnpB functions as a single monomer and its cognate ωRNA shares limited structural similarity to the sgRNA of Cas12f (Supplementary Fig. 3a-d). Moreover, comparisons with other large Cas12 nucleases revealed that the mini-size Cas12f and TnpB interact with nucleic acid less effectively^16, 19–23, 26, 27, 38, 42–45^, which may explain their naturally-low cellular activity and present clues for their rational engineering (Supplementary Fig. 3b-i).

**Figure 1.**
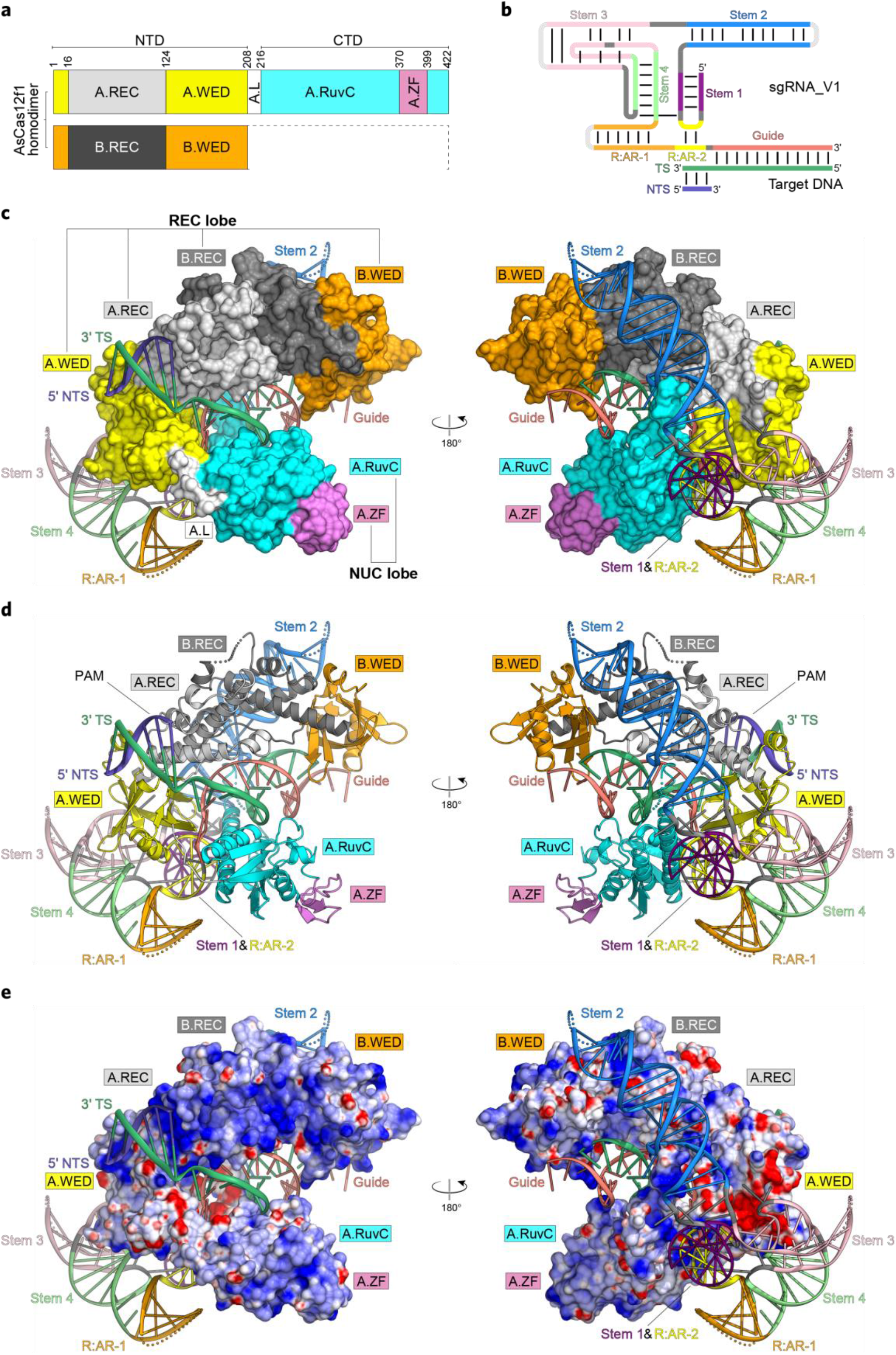
Cryo-EM structure of the AsCas12f1-sgRNA-dsDNA ternary complex. **a**, Domain organization of AsCas12f1. Prefix A and B represent two individual AsCas12f1 monomers in the complex, respectively; NTD, amino-terminal domain; CTD, carboxy-terminal domain; REC, recognition domain; WED, wedge domain; L, linker region; RuvC, RuvC nuclease domain; ZF, zinc-finger domain. The disordered regions are indicated as dashed lines. **b**, Schematic representation of the AsCas12f1 sgRNA architecture. **c**-**e**, The overall structure of the AsCas12f1-sgRNA-dsDNA ternary complex in surface (**c**), ribbon (**d**), and electrostatics (**e**). TS, target strand; NTS, non-target strand; sgRNA, single-guide RNA. The disordered regions are indicated as dotted lines.

### AsCas12f1 effector complex is structurally different from Un1Cas12f1

To understand why AsCas12f1 is an active genome editor with such a compact molecular size that is even smaller than Un1Cas12f1, we comprehensively analyzed the structural differences between the two Cas12f1 nucleases and their corresponding sgRNAs (Fig. 2a-h). Through sequence alignment between AsCas12f1 and Un1Cas12f1, we found that they share similar overall architectures, except that Un1Cas12f1 harbors an exclusive NTD ZF1 domain inserted between the β1 strand of WED and the REC (Supplementary Fig. S4a, b). Next, through structural overlaying, more differences are revealed. As shown in Fig. 2c-e, the B.CTD containing the B.RuvC and B.ZF in AsCas12f1 is completely missing, whereas the counterparts of Un1Cas12f1 can be observed in the structure, except for its B.ZF1 domain in B.NTD, indicating differential molecular flexibilities between the two Cas12f1 nucleases. Furthermore, we found that the dimer interface in AsCas12f1 is larger than that in Un1Cas12f1 (Fig. 2c-e). The exact REC dimer interface area of AsCas12f1 and Un1Cas12f1 are 997.59 Å^2^, and 840.54 Å^2^, respectively. The REC domain of AsCas12f1 comprises five α helixes, while that of Un1Cas12f1 is consisted of four α helixes (Supplementary Fig. S5a). The α1 helix and the loop behind the α5 helix provided the main interactions between the two AsCas12f1 monomers (Fig. 2a, and Supplementary Fig. S5a). Through sequence alignment, we found that the α1 helix in AsCas12f1 is longer than that in Un1Cas12f1 (Supplementary Fig. S4b). Interestingly, the α1 helix (W17-N55) in A.REC is partitioned into two helixes, α1’ (W17-W45) and α1’’ (D51-D54) in B.REC (Supplementary Fig. S5a). The α1’’ helix in B.REC bends towards the α1 helix in A.REC, reinforcing the interactions in the AsCas12f1 dimer, which is discussed in detail below. In contrast, such a difference in A.REC and B.REC is not observed in Un1Cas12f1.

**Figure 2.**
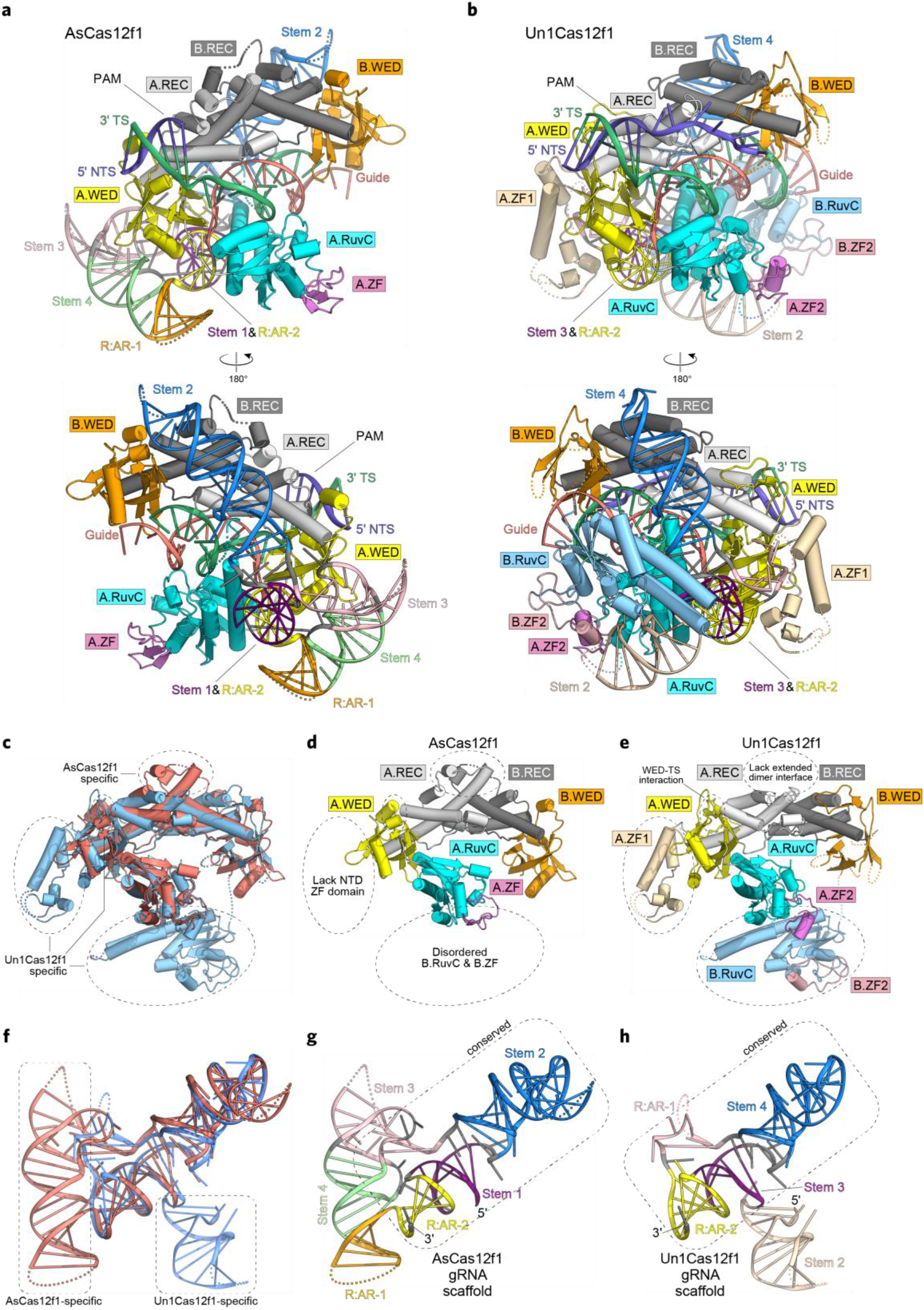
Structural comparison between AsCas12f1 and Un1Cas12f1. **a**, **b**, Structure overview of AsCas12f1 (**a**) and Un1Cas12f1 (**b**). **c**, Nuclease structure overlay of AsCas12f1 (salmon) and Un1Cas12f1 (sky blue). Major structural differences between AsCas12f1 and Un1Cas12f1 are enclosed by dashed ellipses. **d**, **e**, Nuclease structure of AsCas12f1 (**d**) and Un1Cas12f1 (**e**). Structural counterparts are indicated in the same color. **f**, sgRNA structure overlay of AsCas12f1 (salmon) and Un1Cas12f1 (sky blue). Major structural differences between AsCas12f1 and Un1Cas12f1 are enclosed by dashed round square. **g**, **h**, sgRNA structure of AsCas12f1 (**g**) and Un1Cas12f1 (**h**). Structural counterparts are indicated in the same color.

The WED domains in the two Cas12f1 nucleases shared a conserved seven-stranded β-barrel structure core, flanking with different accessories (Supplementary Fig. S5b). In Un1Cas12f1, an extended loop is inserted between the β2 and β3 strands, which provides additional interactions with the TS DNA, narrowing the targeting scope of Un1Cas12f1 (Fig. 2b, and Supplementary Fig. S5b). By contrast, the β2 and β3 strands are linked with the short α1 helix in the REC domain of AsCas12f1. Such additional interactions between WED and TS are not present in AsCas12f1, which is discussed in detail below. Furthermore, Un1Cas12f1 adopts a long β hairpin to connect the β4 and β5 strands, together with the β barrel, forming a hand-like structure to clamp the α1 helix in the REC domain (Supplementary Fig. S5b), while the β4 and β5 strands are connected by the short α2 helix in the REC domain of AsCas12f1. Moreover, the RuvC domains of the two Cas12f1 are highly conserved (Supplementary Fig. S4b, and S5c), and also homologous to other type V CRISPR-Cas nucleases (Supplementary Fig. S3a-i). A ZF domain is inserted into the RuvC domain between its β5 strand and α5 helix in AsCas12f1, containing a β hairpin and a CCCC-type ZF (Supplementary Fig. S5c).

Structural comparisons between the sgRNA scaffolds of AsCas12f1 and Un1Cas12f1 revealed that they share a conserved L-shaped sgRNA core (Fig. 2f-h), which is consisted of the stem 1, 2, partial stem 3, and the R:AR-2 region in AsCas12f1. In contrast, the sgRNA core is formed by the stem 3, 4, partial R:AR-1, and the R:AR-2 region for Un1Cas12f1. Although previous RNA secondary structure predictions indicated that they all shared a similar 5-stem configuration^17, 18, 41^, the actual RNA organizations and the tertiary structures are largely different^19, 20^. Interestingly, the stem 3, 4, and the R:AR-1 region in the AsCas12f1 sgRNA form a large coaxial RNA helix, which is absent in the Un1Cas12f1 sgRNA (Fig. 2f-h). We speculate that the large coaxial RNA helix in AsCas12f1 sgRNA may serve as a surrogate for the NTD ZF1 domain in Un1Cas12f1, whose function is stabilizing the proximal regions of R:AR1 and R:AR2 in the sgRNA core and maintaining the interactions between AsCas12f1 and the sgRNA core. Our previous study on the AsCas12f1 sgRNA indicated that truncations on stem 3 and 4 abolished the dsDNA cleavage activity of AsCas12f1 (ref.^17^), together with the findings in the sgRNA structure, suggesting that the large coaxial RNA helix is indispensable for function. In addition, the stem 2 in the sgRNA of Un1Cas12f1 is located at the 5’ end of the sgRNA core, which interacts with the B.CTD and limits its structural flexibility. In contrast, no RNA component is present at the 5’ end of the sgRNA core in AsCas12f1 sgRNA, and consequently, the B.CTD of AsCas12f1 is flexible in structure. Next, we compared the expression level of protein (HA-tagged) and sgRNA of the two systems in HEK293T cells by western blot and qRT-PCR, respectively (Supplementary Fig. 6a, b). Both the protein expression level and sgRNA transcript level of CRISPR-AsCas12f1 are significantly higher than those of CRISPR-Un1Cas12f1. Similar difference was also observed in a recent study^46^. The difference in protein and sgRNA expression between the two systems may also contribute to their difference in the cellular activities.

Collectively, these data show that AsCas12f1 is structurally different from its ortholog Un1Cas12f1 by its unique dimer configuration and sgRNA architectures, which also explain that AsCas12f1 accepts a large coaxial RNA helix for stabilizing the sgRNA core as a trade-off for the absence of a NTD ZF domain. Therefore, AsCas12f1 (422 a.a.) can utilize even smaller molecular size than Un1Cas12f1 (529 a.a.) for the nuclease functions.

### AsCas12f1 is naturally dimeric through reinforced dimer interface interactions

Two AsCas12f1 monomers predominantly bind with each other to form an asymmetric homodimer through three layers of interactions across their REC domains (Fig. 3a). The bottom layer contains the hydrogen-bond interactions between the loops behind the α5 helix in each REC domain via A.S115 to B.S118, and A.S118 to B.S115 (Fig. 3b). The intermediate layer is mainly formed by the Van der Waals interactions between A.W43, B.W43, and B.I116 (Fig. 3b). Interestingly, the α1’’ helix in B.REC bends towards the α1 helix in A.REC, conferring the additional top-layer dimeric interaction (Fig. 3a). This top-layer dimeric interaction comprises the hydrogen-bond interactions centralized by A.E44 with its surrounding residues, B.W43, B.S50, B.Y52, and B.K53, and the Van der Waals interactions with A.F48 as the core and its surrounding residues, A.W45, A.Y52, B.Y52, and B.H56 (Fig. 3b). Alanine substitutions of S115 and S118 showed no effect on dsDNA cleavage activity (Fig. 3c), whereas the W43A mutation reduced the cleavage activity for ∼13%. In contrast, the top-layer mutations, E44A and F48A, significantly reduced the cleavage activity to ∼51% and ∼22%, respectively. Through size-exclusion chromatography (SEC) analyses, we found that the wild-type AsCas12f1 is naturally a dimeric protein (Fig. 3d). The W43A, F48A, S115A, or S118A mutation can abolish the spontaneous dimerization of AsCas12f1 (Fig. 3d). Although the S115A and S118A mutants are predominantly a monomer in solution, they still have the full nuclease activity (Fig. 3c), suggesting that sgRNA is required for the dimerization process under this circumstance (Supplementary Fig. 7). In addition, the top-layer interactions centralized by E44 and F48 are most important for dimerization and nuclease activity. We speculate that the bending of the α1’’ helix in B.REC towards the α1 helix in A.REC serves as an intermolecular lock, providing additional interactions to stabilize the AsCas12f1 asymmetric homodimer for sgRNA accommodation and target DNA cleavage (Fig. 3a). Therefore, the AsCas12f1-sgRNA ribonucleoprotein (RNP) assembly process can be considered as a binary binding reaction with a stoichiometry ratio at 1:1 for the AsCas12f1 dimer and its cognate sgRNA. By contrast, Un1Cas12f1 is a monomer in solution, whose dimerization is mainly dependent on the binding with sgRNA^19^. Thereby, the stoichiometry ratio for Un1Cas12f1 binding its corresponding sgRNA would be 2:1. Taken together, these findings suggest that the bending of the α1’’ helix in B.REC is a prominent structural feature for maintaining the dimeric configuration of AsCas12f1.

**Figure 3.**
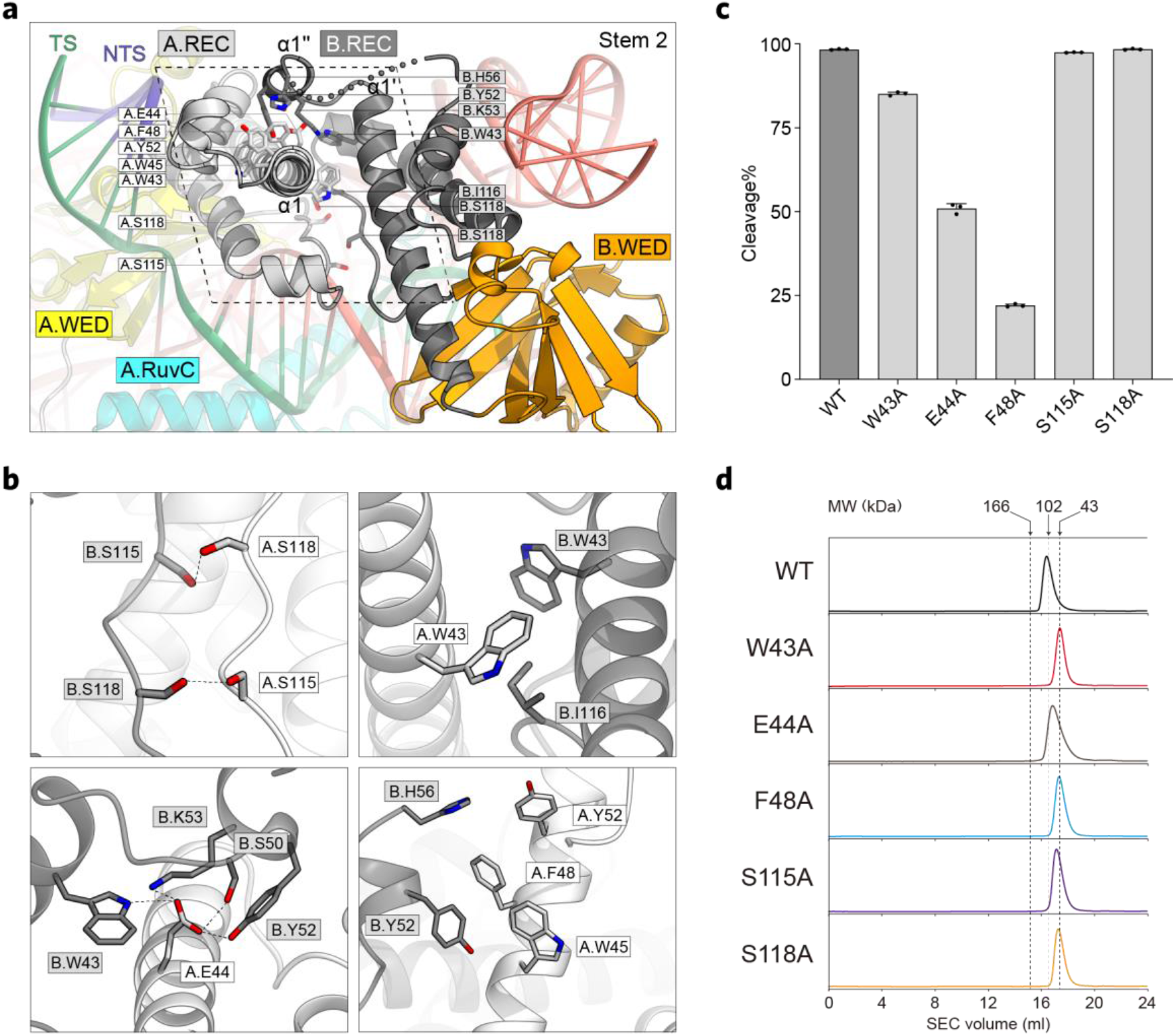
Molecular interactions in the AsCas12f1 homodimer. **a**, Dimer interfaces between AsCas12f1.A and AsCas12f1.B. **b**, Primary interactions between A.REC and B.REC. **c**, In vitro dsDNA cleavage activities of the wild-type AsCas12f1 and dimer-interface mutants. Data represent mean ±SD (n=3). **d**, Size-exclusion chromatography analyses for the wild-type AsCas12f1 and dimer-interface mutants.

### Structural basis for AsCas12f1 PAM preference

AsCas12f1 generally recognizes 5’-NTTR (where R represents A and G) PAMs^17^. To uncover the PAM recognition mechanism, a dsDNA substrate containing a 5’-GTTG PAM was used in structure determination. As shown in Fig. 4a, the PAM duplex is mainly recognized by the A.REC and A.WED domains. Four residues are identified to directly interact with the nucleobases in the PAM duplex. A.Y76 interacts with the nucleobases of dT(-3*) and dT(-2*) in PAM through Van der Waals interactions, and also interacts with the backbone phosphate group of dT(-3*) (Fig. 4b-c). The A.Y76-based interactions are the driving force for the T-rich PAM specificity of AsCas12f1. A.S92 forms hydrogen-bond interactions with the nucleobases of dT(-2*) and dA(-2) (Fig. 4b-c). In addition, A.H72 and A.K96 also interact with the nucleobases of dG(-1*) and dC(-1) through forming hydrogen bonds, respectively (Fig. 4b-c). Alanine substitutions on H72, Y76, S92, or K96 significantly reduced the DNA cleavage activities of AsCas12f1 (Fig. 4d), suggesting that these four residues are critical for the PAM recognition by AsCas12f1. In addition, A.K63, A.Y68, A.T69, A.K80, A.S88, A.K129, A.N151, A.S147, and A.K154 extensively interact with the backbone phosphate groups of the PAM duplex through forming hydrogen bonds (Fig. 4b). By contrast, Un1Cas12f1 typically specifies 5’-TTTR PAMs^18–20, 36, 37^. A.Y146 of Un1Cas12f1 is functional equivalent to A.Y76 of AsCas12f1, together with A.A156, interacting with the nucleobases of dT(-2*), dT(-3*), and dT(-4*) in PAM through Van der Waals interactions (Fig. 4e)^19^. A.S142 and A.R163 interact with the nucleobases of dG(-1*) and dC(-1) through forming hydrogen bonds (Fig. 4e). In addition, A.Q197 and A.Y202 located on the extended β2-to-β3 loop in the A.WED domain of Un1Cas12f1 provides additional hydrogen-bond interactions with dA(-3) and dA(-4), respectively (Fig. 4e, and Supplementary Fig. S5b). Consequently, these structural configurations limit Un1Cas12f1 to recognize 5’-TTTR PAMs. By contrast, the β2 and β3 strands are connected by a short α helix in the REC domain of AsCas12f1 (Supplementary Fig. S5b), and thereby, the A.WED domain marginally interacts with the TS-side DNA in the PAM duplex (Fig. 4a-c), explaining why AsCas12f1 has broader targeting scopes than Un1Cas12f1.

**Figure 4.**
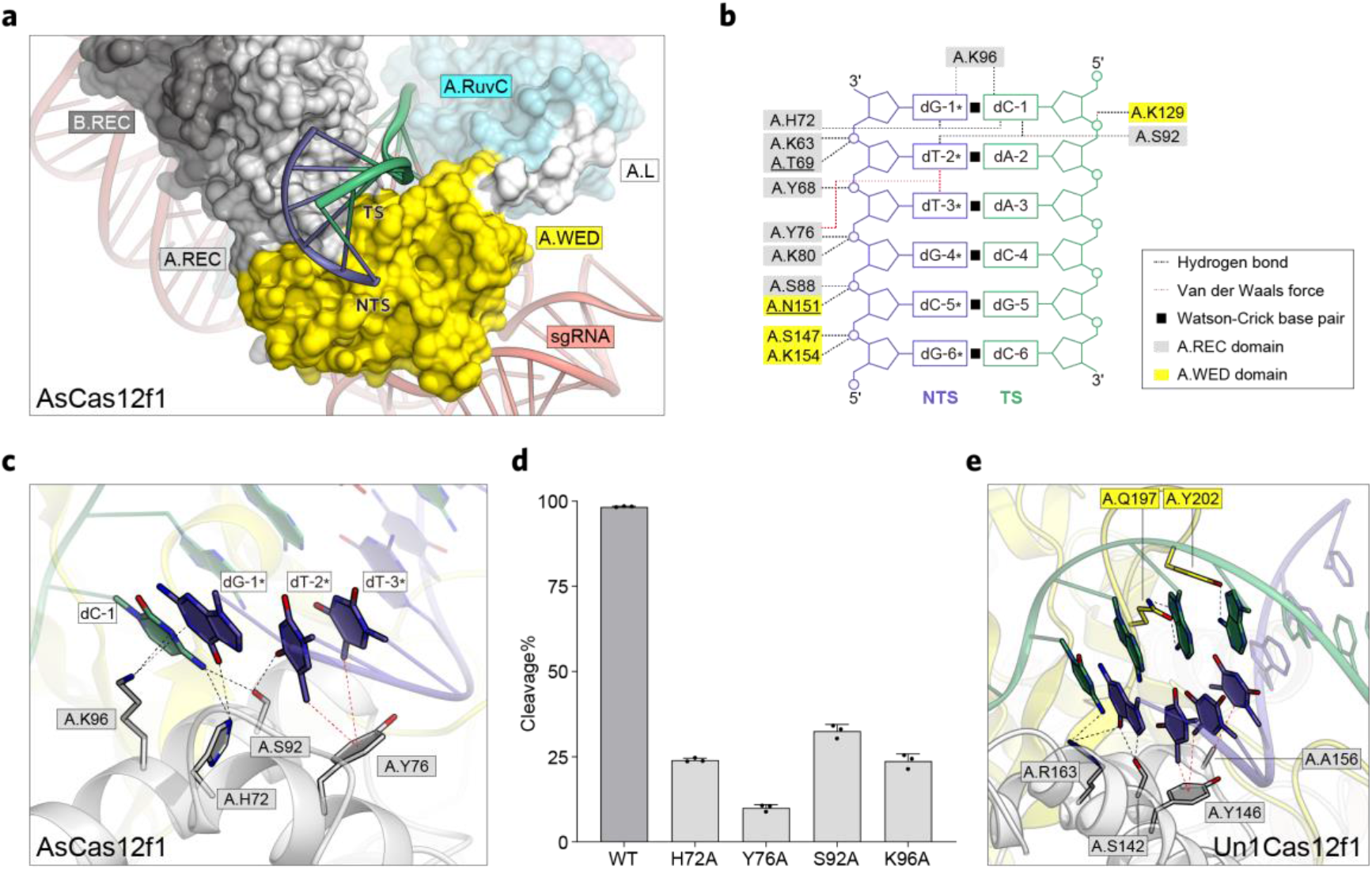
PAM recognition mechanism. **a**, PAM duplex binds to the groove between A.REC and A.WED domain. **b**, Molecular interactions between AsCas12f1 and nucleotides in the PAM duplex. **c**, Key interactions between bases in the PAM duplex and PAM-recognizing residues of AsCas12f1. **d**, In vitro dsDNA cleavage activities of the wild-type AsCas12f1 and PAM-recognizing mutants. Data represent mean ±SD (n=3). **e**, Molecular interactions in PAM recognition by Un1Cas12f1.

The heteroduplex of the target DNA and the sgRNA is accommodated within the positively charged central groove of the AsCas12f1 dimer through sequence-independent interactions against the sugar-phosphate backbones (Fig. 1e, and Supplementary Fig. S8a-e). A.K3, A.Y351, A.H303, A.R6, A.N204, A.K120, A.Q191, A.R101, A.S118, and A.S115 extensively interact with the sugar-phosphate backbone of the first seven ribonucleotides in the sgRNA (C172 to U179) (Supplementary Fig. S8a, b). A.Y343, A.R298, A.R291, and A.R280 interacts with the sugar-phosphate backbone of dC(6), dA(7), dA(8), and dA(11) in TS (Supplementary Fig. S8a, b). However, the map density does not support the model of the structure for nucleotides 14-24 of the TS and 189-191 of the sgRNA.

Collectively, the structural knowledge provided here may contribute to the nuclease engineering for enhancing the genome editing activity of AsCas12f1.

### Unique large coaxial RNA helix confers AsCas12f1 sgRNA extra structural stability

AsCas12f1 essentially requires a tracrRNA and a crRNA for DNA cleavage. To simplify the delivery, the two RNAs are joined together with a tetraloop, generating the sgRNA_V1 (191 nt) with enhanced DNA cleavage activity (Fig. 5a-b)^17^. This initial version of sgRNA is sufficient to drive the AsCas12f1-mediated genome editing in bacteria and mammalian cells^17^. In comparison, the Un1Cas12f1-sgRNA system showed no plasmid-interference activity in *E. coli* and marginal genome-editing activities in mammalian cells^33, 41^. Through removing the structurally flexible RNA components from the sgRNA, the potential of Un1Cas12f1 as a genome editor was eventually unleashed^19, 20, 36, 37^, suggesting that the structural stability of sgRNA determines the in vivo nuclease activity of Cas12f1. Structural comparisons between the effector complexes of AsCas12f1 and Un1Cas12f1 reveals a conserved sgRNA core, comprising the stem 1, 2, partial stem 3, and the R:AR-2 region (Fig. 2f-h), which extensively interacts with the AsCas12f1 dimer (Supplementary Fig. S8a-e). The long α-helix hairpin formed by the α1 and α2 helixes in A.RuvC interposes into the groove (G27-G32 and C63-C67) of stem 2. The ribonucleotides G29-A31-C65 and A30-G32-C63 form two layers of base triplexes, respectively, and G64 flips out from the stem 2 (Fig. 5a-c, and Supplementary Fig. S8a). Similar local sgRNA configurations were also observed in the stem 4 of Un1Cas12f1 sgRNA^19, 20^. The terminal of the stem 2 interacts with the PAM-recognizing residues in B.REC, however, through a sequence-independent manner by forming hydrogen bonds with the sugar-phosphate backbone (Supplementary Fig. S6a, d). The R:AR-2 duplex together with the stem 1 forms a coaxial RNA helix, which extensively interacts with the A.WED and A.RuvC domains (Supplementary Fig. S8a, e). Besides the sgRNA core, the large coaxial RNA helix formed by the stem 3, 4, and the R:AR-1 duplex significantly contributes to the structural stability of the AsCas12f1 sgRNA. The ribonucleotides G74-A90-C107, U75-A89-C101-A104, and A76-G88-A103 form three layers of base triplexes and quadruplex (Fig. 5a-d, and Supplementary Fig. 8a). G105 and A106 flip out from the stem 3, and interact with A.W17 and A.D16, respectively. G109, C110, and A111 in stem 4, interact with A.H139, A.R136, and A.K147, respectively. Moreover, C92 interacts with G14 via canonical Watson-Crick base pairing (Supplementary Fig. 8a). The large coaxial RNA helix utilizes these unique interactions to stabilize the binding of AsCas12f1 to the sgRNA core, and takes the place of the NTD ZF1 domain in Un1Cas12f1, allowing AsCas12f1 to serve as an active genome editor with such a compact molecular size (422 a.a.) that is even ∼100 a.a. smaller than Un1Cas12f1 (529 a.a.).

**Figure 5.**
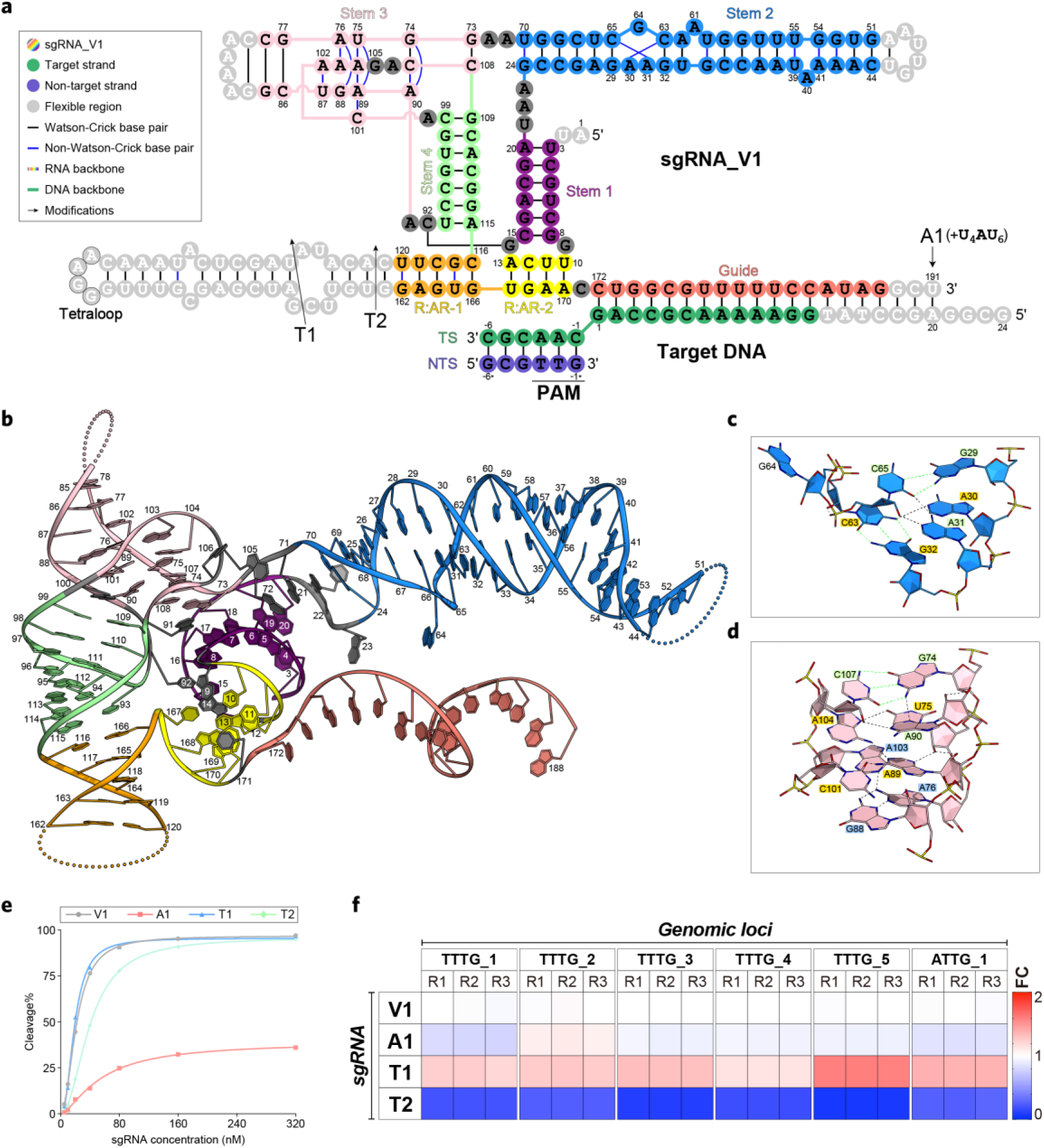
Architecture of the single-guide RNA and its engineering. **a**, Schematic representation of the sgRNA and the target DNA. **b**, The structure of the sgRNA. **c**, Base triplexes in the Stem 2 of the sgRNA scaffold. **d**, Base triplexes and quadplex in the Stem 3 of the sgRNA scaffold. **e**, Dose-response in vitro dsDNA cleavage activities of the wild-type AsCas12f1 with different modifications of sgRNA. Scatters represent mean of triplicates. **f**, Indel activities of the wild-type AsCas12f1 (relative to sgRNA_V1) with different modifications of sgRNA on 6 genomic loci in HEK293T cells, determined by NGS.

Previous studies indicated that the complete stem 1, the distal region of stem 2, and R:AR-1 of the Un1Cas12f1 sgRNA were flexible in the complex structures^19, 20^. Truncations in these regions result in an engineered version of sgRNA with enhanced in vivo nuclease activity of Un1Cas12f1^37^. However, our previous study showed that only the long-extended RNA duplex in the distal region of R:AR-1 can be truncated without adverse effect on DNA cleavage activity^17^, which is in agreement with the findings in the AsCas12f1 sgRNA structure, as the distal region of R:AR-1 is flexible without interactions with AsCas12f1 and the sgRNA core (Fig. 5a, b). Intrigued by the engineering of the Un1Cas12f1 sgRNA^37^, we sought to investigate whether the removal of the redundant RNA components in AsCas121 sgRNA, and the addition of polyU (U_4_AU_6_) to the 3’-terminal of the sgRNA, can contribute to the improved dsDNA-cleaving and genome-editing activities (Fig. 5a). The sgRNA-dose-dependent dsDNA cleavage assays showed that the T1 truncation exhibited similar dsDNA-cleaving activity to the original sgRNA_V1, while the activity slightly reduced in the T2 truncation (Fig. 5e), indicating the active boundary for the R:AR-1 region. Moreover, the dsDNA-cleaving activity is significantly impaired by the A1 addition (Fig. 5e), suggesting that adding polyU to the 3’-terminal of the guide RNA introduces mismatches behind the guide RNA-target DNA duplex and undermines the dsDNA cleavage process. Next, we explored the effect of sgRNA modifications on cellular indel activity. As shown in Fig. 5f, the T1 truncation elevated the indel activities of AsCas12f1 on 6 genomic loci in HEK293T cells. The indel activity was significantly attenuated by the T2 truncation, again confirming the active boundary for the R:AR-1 region. However, the A1 addition showed comparable or slightly lower indel activities than the original sgRNA_V1 (Fig. 5f). This observation is opposite to the circumstance for Un1Cas12f1 sgRNA engineering, emphasizing the difference between the two systems. The addition of polyU (U_4_AU_6_) to the 3’-terminal of the sgRNA was originally proposed for improving the in vivo stability of the Un1Cas12f1 sgRNA^37^. In contrast, the AsCas12f1 sgRNA_V1 is sufficiently stable for in vivo genome editing, and thereby, the polyU stabilizer may not be necessary for the AsCas12f1 sgRNA. Together, an engineered sgRNA_T1 was obtained with enhanced genome-editing activity in mammalian cells in a shortened sgRNA length (162 nt).

### Molecular mechanism for AsCas12f1-mediated dsDNA cleavage

Most Cas12 family nucleases harbor a single RuvC nuclease domain for the cleavage of both the TS and NTS^2^. However, Cas12f1 nucleases form a dimer in the effector complex, presenting two RuvC nuclease domains. Previous studies on Un1Cas12f1 revealed that only the A.RuvC was responsible for target DNA cleavage, while the nuclease-active site of B.RuvC was occupied by the two flipped ribonucleotides of the stem 2 (ref.^19, 20^). By contrast, the AsCas12f1 sgRNA does not have such a RNA counterpart to contact with the B.RuvC (Fig. 2f-h), and therefore, the complete B.CTD of AsCas12f1 is flexible in the effector complex (Fig. 1c-e).

To determine the key residues for the nuclease activity of AsCas12f1, alanine substitutions were created on D225, E324, D325, N328, D331, R335, D364, S369, R383, Q386, and D401, respectively (Supplementary Fig. 9a, b). Through DNA cleavage assays, we found that D225A, E324A, R383A, and D401A abolished the nuclease activity of AsCas12f1 as previously described (Supplementary Fig. 9c)^17^. R335A and S369A also significantly reduced the nuclease activity. Moreover, to determine whether the flexible B.RuvC domain is involved in dsDNA cleavage, we created covalent dimer variants and mutated the active sites in each monomer as previously described (Supplementary Fig. 10a)^19^. The dsDNA-cleavage activities of these dimer variants were compared with that of wild-type AsCas12f1, showing that only A.RuvC is involved in DNA cleavage (Supplementary Fig. 10b), consistent with the findings in Un1Cas12f1^19^. In addition, the ZF domain nearby the RuvC is proposed to load the TS and NTS into the RuvC active site, like other Cas12 nucleases^2^. Alanine substitutions of C372, C375, C391, and C393, lead to severe protein precipitations. As the map density in this region is not clear enough to identify the presence of Zn ion, we performed the inductively coupled plasma-optical emission spectroscopy (ICP-OES) assay and confirmed that the ZF domain in AsCas12f1 can naturally bind with Zn ion (Supplementary Fig. 11a, b).

### Rational engineering of AsCas12f1 to improve genome-editing activity in mammalian cells

Through comprehensive structural analyses of the AsCas12f1 effector complex, we attributed the naturally-moderate cellular activity of AsCas12f1 to the limited interactions between nuclease and nucleic acid. To improve the genome-editing activity of AsCas12f1, we selected amino acid residues located adjacent to nucleic acids and introduced R, K, and Q substitutions, in order to enhance the binding of AsCas12f1 to sgRNA or DNA. Here, we created 92 residue substitutions in the REC, WED, and RuvC domains (Fig. 6a). The genome-editing activity of these effector variants were directly screened by indel assays at two TTTG-PAM target sites in HEK293T cells. Ten substitutions, E8R, T69S, N70Q, K103R, A104R, K107R, E108R, S115R, S118A, and D364R, were identified with enhanced indel activities (Fig. 6a). These variants were then tested at 6 other target sites, including 2 CTTG-PAM sites, 2 ATTG-PAM sites, and 2 TTTA-PAM sites. E8R and T69S exhibited significantly attenuated indel activities on at least three target sites (Fig. 6b), and thus, they were excluded for the subsequent mutation combination. Next, A104R and E108R with the highest indel activities on the 2 TTTG-PAM target sites, were selected as the starting nucleases for combining other mutations. We found that E108R was incompatible with other mutations, whereas A104R exhibited much higher indel activities in collaboration with other mutations (Fig. 6c, and Supplementary Fig. 12). Finally, a combined variant v5.1 (N70Q+K103R+A104R+S118A+D364R) exhibited highest indel activities across 8 tested target sites, with 1.5∼13.5-fold improvements relative to the wild-type AsCas12f1 (Fig. 6c).

**Figure 6.**
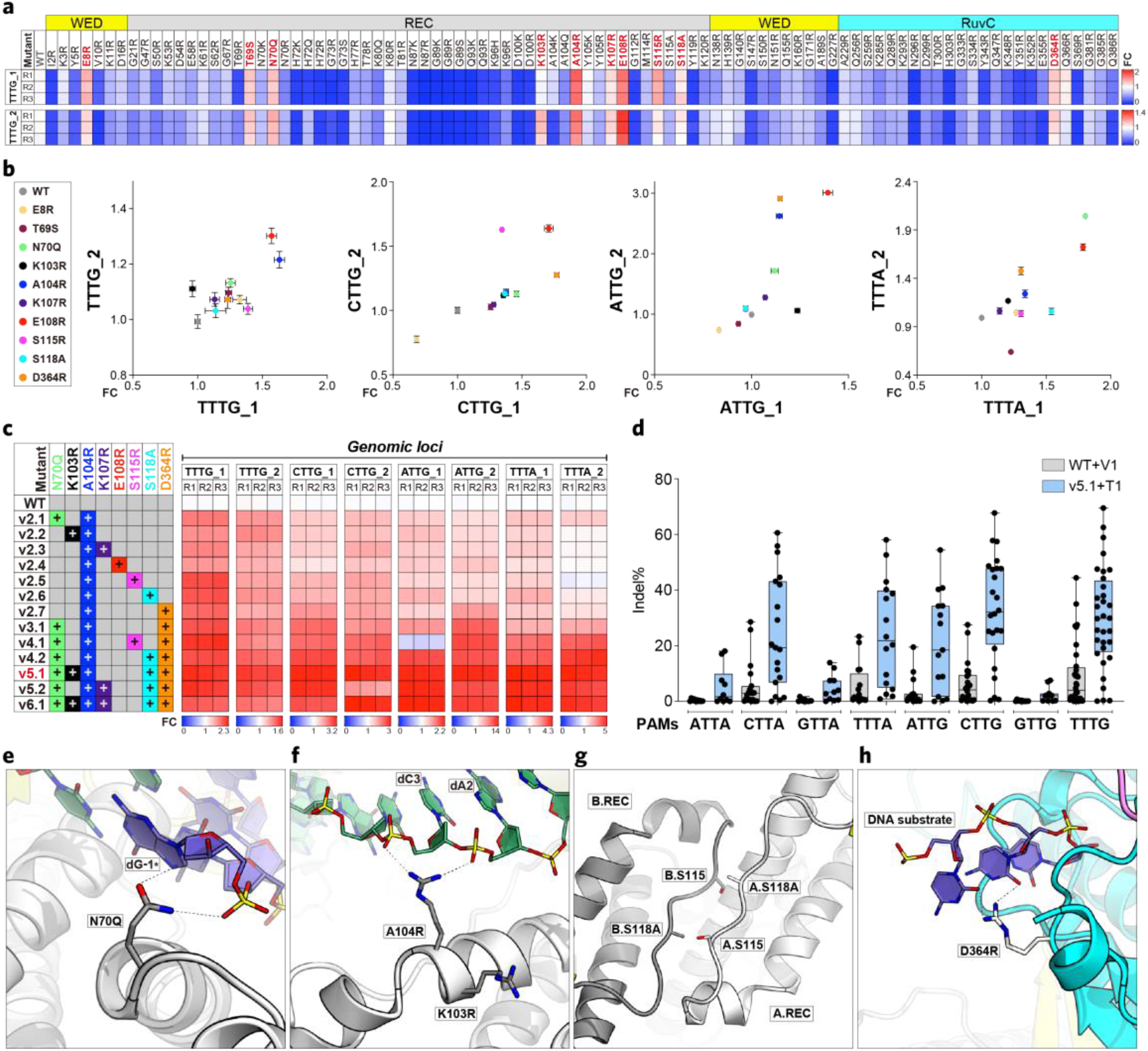
Structure-directed engineering of AsCas12f1. **a**, Direct mutation screen of genome-editing activity (relative to WT) on two TTTG-PAM genomic loci in HEK293T cells, determined by NGS. **b**, Indel activity (relative to WT) of 10 single-substitution variants measured in HEK293T cells at 8 genomic loci, determined by NGS. Data represent mean ± SD (n=3). **c**, Indel activity (relative to WT) of the combination variants, v2.1∼2.7, v4.1, v4.2, v5.1, v5.2, and v6.1, measured in HEK293T cells at 8 genomic loci, determined by NGS. **d**, Comparison of indel activity between the engineered AsCas12f1-v5.1+T1 and the wild-type AsCas12f1+V1 in HEK293T cells at 138 genomic loci covering all 8 types of NTTR PAMs, determined by NGS. Scatters represent mean of triplicates. Center line represents the median. Top and bottom lines represent the upper and lower extremes. Box represents the upper and lower quartiles. **e**, **f**, **g**, **h**, The structure basis of the activity-enhancing mutations, N70Q (**e**), K103R, A104R (**f**), S118A (**g**), and D364R (**h**).

To comprehensively assess the genome-editing activity of the engineered AsCas12f1-v5.1+T1, we tested the indel activities on 138 genomic loci covering all 8 types of NTTR PAMs in comparison with the wild-type AsCas12f1+V1. As shown in Fig. 6d, the overall indel activity of AsCas12f1 was significantly elevated through the structural-guided rational design. The indel activities of the engineered AsCas12f1-v5.1+T1 were maximal on TTTG- and CTTG-PAM target sites, moderate on CTTA-, TTTA-, and ATTG-PAM target sites, and minimal on ATTA-, GTTA-, and GTTG-PAM target sites (Fig. 6d). Moreover, the indel activity of the engineered AsCas12f1-v5.1+T1 were also tested in Hela cells (Supplementary Fig. 13).

Structural analyses indicate that the N70Q mutation fortifies the interaction between the A.REC domain and dG(-1*) in the PAM duplex (Fig. 6e). The A104R mutation provides new interactions between the A.REC domain and the sugar-phosphate backbone of dA(2) and dC(3) in TS, while the K103R mutation may stabilize the displaced NTS (Fig. 6f). Intriguingly, the monomeric mutation S118A enhanced the indel activity of AsCas12f1, indicating that the dimerization of AsCas12f1 is not necessary for its cellular activity (Fig. 6a-c, g). Moreover, the D364R mutation is supposed to reinforce the interaction of A.RuvC with the DNA substrate (Fig. 6h).

To provide a benchmark, we systematically compared the genome-editing activity of the engineered AsCas12f1-v5.1+T1 with Un1Cas12f1-v3.1+ge4.1, enAsCas12a, and SpCas9 across 14 different genomic loci. These target sites concurrently have 5’ TTTG or TTTA PAMs and 3’ NGG PAMs in order to suit the 4 Cas nucleases in the comparison. The results showed that the engineered AsCas12f1-v5.1+T1 exhibits comparable editing activity with enAsCas12a and SpCas9, and higher activity than Un1Cas12f1-v3.1+ge4.1, at most of the tested loci (Fig. 7a). Next, the genome-wide off-target editing activity of the engineered AsCas12f1-v5.1+T1 was comprehensively compared with Un1Cas12f1-v3.1+ge4.1, enAsCas12a, and SpCas9 across 6 different genomic loci using the widely-adopted off-target profiling assay GUIDE-seq^47^. As shown in Fig. 7b-h, two engineered Cas12f1 nucleases exhibited much lower off-target editing activities than enAsCas12a and SpCas9 at most of the tested sites, as reflected by the proportion of off-target editing events in total editing events. This finding was also supported by other recent comparative studies^46, 48^. Moreover, AsCas12f1-v5.1+T1 showed similar or lower off-target editing activities than Un1Cas12f1-v3.1+ge4.1 at 4 target sites, TTTG_1, TTTA_2, TTTA_18, and TTTA_21 (Fig. 7c, f-h). Although Un1Cas12f1-v3.1+ge4.1 showed no off-target event at 3 TTTG sites, their on-target read numbers were substantially lower than these observed on AsCas12f1-v5.1+T1 (Fig. 7c-e), indicating that the editing activity of Un1Cas12f1-v3.1+ge4.1 was lower than that of AsCas12f1-v5.1+T1 at these editing loci, which may underestimate the off-target effect caused by Un1Cas12f1-v3.1+ge4.1.

**Figure 7.**
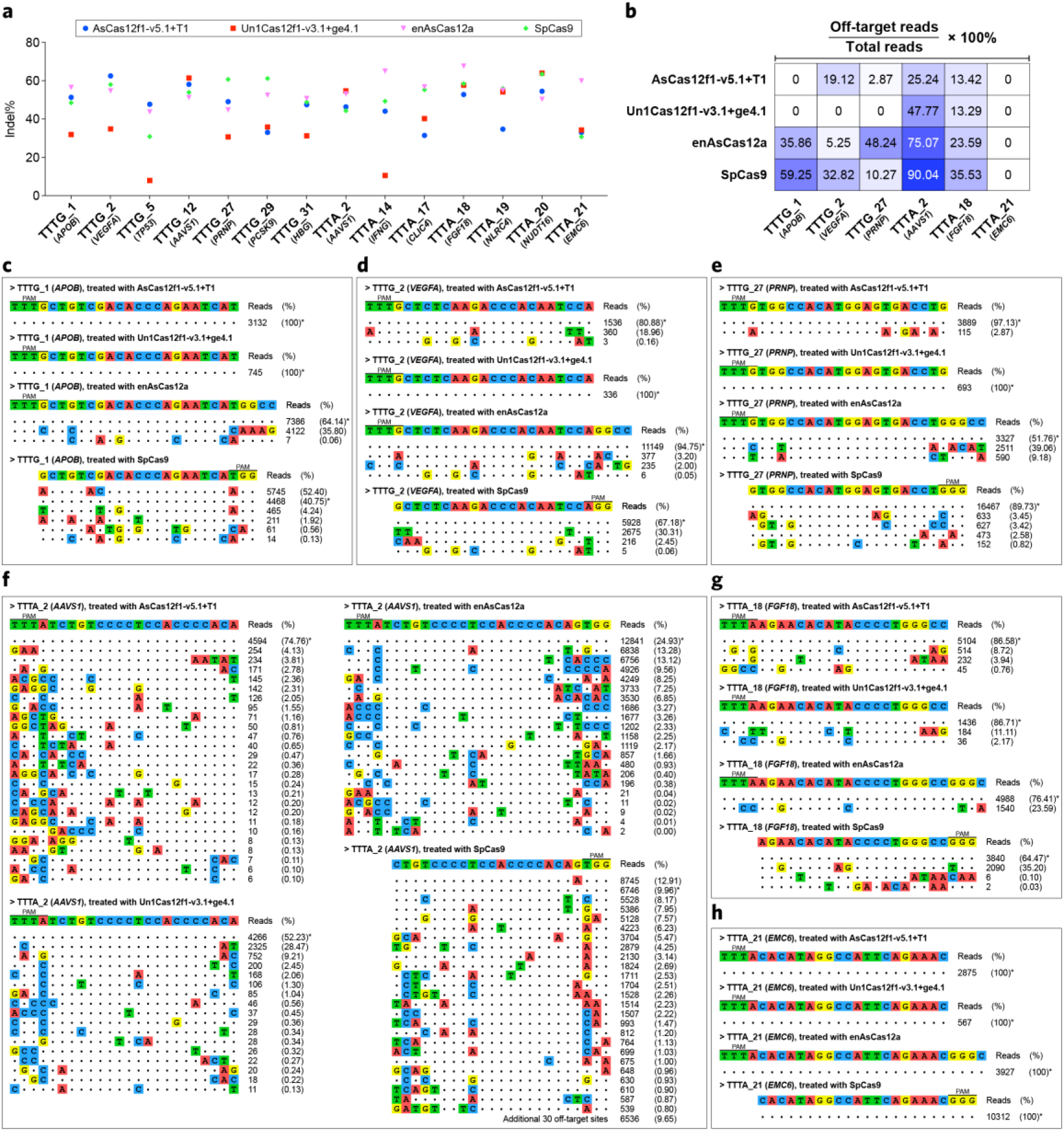
Systematic evaluation of the engineered CRISPR-AsCas12f1 system. **a**, Comparison on indel activities of AsCas12f1-v5.1+T1 with Un1Cas12f1-v3.1+ge4.1, enAsCas12a, and SpCas9 across 14 different genomic loci in HEK293T cells, determined by NGS. Scatters represent mean of triplicates. **b**, Comparison on the proportion of off-target editing events in total editing events across the four tested genome editor, determined by GUIDE-seq. **c**, **d**, **e**, **f**, **g**, **h**, Off-target editing events caused by AsCas12f1-v5.1+T1 with Un1Cas12f1-v3.1+ge4.1, enAsCas12a, and SpCas9 in HEK293T cells at TTTG_1 (**c**), TTTG_2 (**d**), TTTG_27 (**e**), TTTA_2 (**f**), TTTA_18 (**g**), and TTTA_21 (**h**), respectively.

Taken together, these data demonstrate that the engineered AsCas12f1-v5.1+T1 is an efficient and specific genome editor in mammalian cells. Given the ultra-compact molecular sizes of AsCas12f1-v5.1 (422 a.a.) and its corresponding sgRNA_T1 (162 nt), this genome-editing tool may serve as a promising candidate for therapeutic applications.

## Discussion

In this study, we demonstrate that the ultra-compact CRISPR-Cas nuclease, AsCas12f1 (422 a.a.), forms an asymmetric dimer to accommodate one sgRNA molecule for target dsDNA recognition and cleavage. This finding explains why AsCas12f1 can function as a programmable CRISPR-Cas nuclease similar to other large Cas12 nucleases with such a small molecular size. In one functional effector unit, two AsCas12f1 monomers cooperate with each other to form a multi-protein, rather than a single-protein effector complex. This finding also challenges the definition for class 2 CRISPR-Cas systems which features in a single large effector nucleases^1^. Moreover, comparative structural studies on the effector complexes of AsCas12f1 and Un1Cas12f1 present clues for understanding the evolution of CRISPR-Cas system and facilitating the engineering of mini-size Cas nuclease.

The two AsCas12f1 monomers in the effector complex vary in their overall and local confirmations, as well as their functions in sgRNA binding, target DNA recognition, and cleavage. Two AsCas12f1 monomers tightly interact with each other through their REC domains. Structural information indicates that the bending of the α1’’ helix in B.REC towards the α1 helix in A.REC provides additional dimerization interactions, given that AsCas12f1 naturally maintains dimeric in solution even in the absence of sgRNA, whereas the dimerization of Un1Cas12f1 is dependent on the addition of sgRNA^19^. Intriguingly, certain monomeric mutations, S115A and S118A, change the manner of dimerization into a sgRNA-dependent mechanism like that of Un1Cas12f1. Nevertheless, these variants still retain the full dsDNA-cleavage activities in comparison to the wild-type AsCas12f1, indicating that the naturally-dimeric configuration of AsCas12f1 is not necessary for its catalytic activity. Moreover, S118A even enhanced the indel activity of AsCas12f1 in human cells.

A unique large coaxial RNA helix formed by stem 3, 4, and R:AR-1, was identified in the AsCas12f1 sgRNA but not present in Un1Cas12f1 sgRNA. The large coaxial RNA helix fortifies the overall stability of AsCas12f1 sgRNA and reinforces the interactions between the sgRNA core and the AsCas12f1 dimer. Through structural comparisons of AsCas12f1 with other Cas12 nucleases, we found that the effector configuration of AsCas12f1 is more analogous to that of Cas12b and Cas12e, as the A.REC, A.WED, B.REC, and B.WED domains join together to form a relatively small REC lobe for PAM recognition and sgRNA binding, while a single A.RuvC and A.ZF construct a small NUC lobe for target DNA cleavage. We speculate that some large Cas12 nucleases are evolved from precursors like AsCas12f1, as they gradually obtained additional function domains and abandoned excess RNA components over time. This idea is reinforced by the structural evidences from the comparison between the effector complexes of AsCas12f1 and Un1Cas12f1. The large coaxial RNA helix in AsCas12f1 sgRNA supersedes the position of the NTD ZF1 domain in Un1Cas12f1 to stabilize the sgRNA core, suggesting that AsCas12f1 may also be a precursor for Un1Cas12f1, which somehow acquired the NTD ZF1 domain and inserted before the REC domain, thereby abandoning some RNA components around its sgRNA core.

The structural information provided in this study also allows us to rationally engineer the CRISPR-AsCas12f1 system to improve its genome-editing activity. Here, an engineered version, AsCas12f1-v5.1 with sgRNA_T1 (v5.1+T1), containing substitutions of N70Q, K103R, A104R, S118A, and D364R, was obtained with drastically enhanced indel activity in human cells. These substitutions likely strengthen the electrostatic interaction with DNA. Such a compact, efficient, and precise genome editor offers particular advantages for easier and more flexible cellular delivery using cargo-size-limited vectors, such as AAV, whereas the Cas9- or Cas12a-based genome editors are usually overweight. Therefore, the future engineering of AsCas12f1-based genome editors may focus on expanding its target scope by fine tuning the PAM-interacting residues, and further enhancing its editing activity by directed evolution, which possibly brings new solutions for some challenges in therapeutic applications.

## Methods

### Protein and RNA preparation

The wild-type AsCas12f1 nuclease and its mutants were prepared as previously described^17^. Briefly, the plasmid encoding the N-terminally His_6_-SUMO-tagged AsCas12f1 was introduced into *E. coli* BL21(DE3) for heterologous protein expression. The *E. coli* cells were cultured at 37°C with vigorously shaking until the OD_600_ reached 0.6, and the protein expression was then induced by adding isopropyl-β-D-thiogalactopyranoside (IPTG) to a final concentration of 0.25 mM, and continued incubating at 16°C for overnight. The *E. coli* cells were pelleted by centrifugation and resuspended in buffer A (20 mM Tris-HCl, pH=7.5, 1 M NaCl, 15 mM imidazole, and 1 mM DTT), and homogenized by using a high-pressure cell disruptor at 900 bar for 5 min. The clarified supernatant was loaded to a pre-equilibrated 5-ml HisTrap Ni-NTA column (Cytiva). Unbound proteins were washed away with buffer A containing 50 mM imidazole, and AsCas12f1 was eluted with buffer A containing 250 mM imidazole. Then, the His_6_-SUMO tag was cleaved by using HRV3c treatment at 4°C for overnight. The digested product was then loaded to a pre-equilibrated HiLoad 16/600 Superdex 200pg column (Cytiva) for size-exclusion chromatography. The purified proteins were stored at -80°C until use. The whole procedure of protein purification was carried out at 4°C. The sgRNAs were transcribed in vitro by using a HiScribe T7 High Yield RNA Synthesis Kit (NEB), and purified by using TRIzol Reagent (Invitrogen) following the manufacturers’ instructions, respectively. Plasmid and sgRNA used in this study are listed in Supplementary Dataset 1.

### Electron microscopy sample preparation

The AsCas12f1-sgRNA-dsDNA complex was assembled by mixing the purified AsCas12f1-D225A mutant, the 191-nt sgRNA_V1, and the target DNA pre-annealed with a 30-nt target strand and a 6-nt non-target strand, at a molar ration of 1:1.5:1.5 in SEC buffer (20 mM Tris-HCl, pH=7.5, 600 mM NaCl, 5 mM MgCl_2_, and 1 mM DTT), and incubate at 45°C for 10 min. The AsCas12f1-sgRNA-dsDNA complex was then purified by size-exclusion chromatography using a pre-equilibrated Superose 6 10/300 GL column (Cytiva). The purified complex was concentrated to approximately 5 mg/ml. Holey carbon grids (Quantifoil Au 300 mesh, R1.2/1.3) were glow-discharged in the Plasma Cleaner PDC-32G-2 (Harrick) with a vacuum for 2 min and mid force for 28 s. Aliquots (3.8 µl) of the complex were placed on the freshly glow-discharged grids, which were then blotted for 3 s and plunged frozen in liquid ethane cooled by liquid nitrogen using Vitrobot Mark IV (ThermoFisher Scientific) at 8℃ and 100% humidity.

### Electron microscopy data collection and processing

Electron micrographs were acquired on a Titan Krios electron microscope (ThermoFisher Scientific) operating at 300 kV and equipped with Cs corrector, Gatan K3 Summit detector, and GIF Quantum energy filter with a slit width of 20 eV. Images were automatically collected using AutoEMation^49^ and a preset defocus range from -1.8 µm to -2.2 µm in super-resolution mode at a nominal magnification of 81,000×. Each stack was exposed for 2.56 s with an exposure time of 0.08 s per frame, resulting in a total of 32 frames per stack and the total dose was approximately 50 e-/Å^2^ for each stack. The stacks were motion corrected with MotionCor2^50^ and binned 2 folds, resulting in a pixel size of 1.0773 Å/pixel. Meanwhile, dose weighting was performed^51^. The defocus values were estimated with Gctf^52^. A diagram for data processing is presented in Supplementary Figure S1. A total of 4,683,726 particles were autopicked with Gautomatch from 4,160 collected micrographs. The autopicked particles were imported to cryoSPARC^53, 54^, wherein 3,407,245 selected particles from several rounds of 2D classifications were subjected to heterogeneous refinement. Then, 1,770,878 particles from the best class were subjected to non-uniform refinement, resulting in a reconstruction of 2.77-Å overall resolution^55^. Next, local CTF refinement and non-uniform refinement improve the final density map to a resolution of 2.69 Å. The resolution was estimated with the gold-standard Fourier shell correlation 0.143 criterion with high-resolution noise substitution^54, 56^.

### Model building and validation

The structure model for AsCas12f1-sgRNA-dsDNA ternary complex was built manually in COOT^57^. The *phenix.real_space_refine* tool^58^ was used to refine the model against the density map. Molecular visualization figures were generated by using CueMol (http://www.cuemol.org).

### Phylogenetic analysis

The sequences of RNA-guided endonuclease TnpB family protein were accessed from NCBI Protein Family Models (accession: NF040570.1). The sequences of type V Cas nucleases were collected from previous studies and BLAST hits from RefSeq^1, 25^. The multiple sequence alignment of the selected proteins was generated by using MAFFT with default parameter setting^59^. The phylogenetic tree was constructed using FastTree (JTT+CAT)^60^, and visualized in iTOL using TnpB as the root of the phylogenetic tree^61^.

### In vitro cleavage assay

The dsDNA cleavage activities of the wild-type AsCas12f1 and its mutants were determined as previously described with minor modifications^17^. Briefly, 5 nM of linear dsDNA substrates were mixed with 150 nM of AsCas12f1 and 300 nM of sgRNA in 1× cleavage buffer (10 mM Tris-HCl, pH=7.5, 10 mM MgCl_2_, and 50 mM NaCl), and incubated at 45°C. The cleavage reactions were stopped at 32 min by adding the quench buffer (50 mM EDTA, 1 mg/ml RNase A, and 1 mg/ml Proteinase K). The digest products were further incubated at 37°C for 10 min and 50°C for 10 min to degrade sgRNA and nuclease, and then analyzed by agarose gel electrophoresis.

### Inductively coupled plasma-optical emission spectroscopy (ICP-OES)

To examine the Zn ion content within AsCas12f1, ICP-OES assays were carried out as previously described with modifications^62^. Briefly, the purified AsCas12f1 protein was air dried at 95°C for overnight, and then digested by 500 μl of nitric acid (66-68%) for 6 h at 80°C. The digest was diluted with ddH_2_O to 10 ml, and subjected to ICP-OES assays using iCAP 7400 (ThermoFisher Scientific). Standard Zn ion solutions were also examined in parallel.

### Cell culture and transfection

The HEK293T cells were routinely maintained in DMEM (Corning) with 10% fetal bovine serum (ExCell) at 37°C in 5% CO_2_, and split every 48 h to ensure the cells never exceed 95% confluency. For plasmid transfection, cells were seeded into 24-well plates at 0.8×10^5^/well. After 16-h incubation, plasmid was transfected by using Lipofectamine 3000 (Invitrogen) according to the manufacturer’s instructions. The cells were harvested 48-h post transfection to evaluate the genome editing efficiency.

### Expression profiling

To determine the expression level of AsCas12f1 and Un1Cas12f1 in mammalian cells, the C-terminal HA-tagged Cas12f/sgRNA-encoding plasmids were transfected into HEK293T cells by using Lipofectamine 3000 (Invitrogen) according to the manufacturer’s instructions. Cells were harvested 48 h post transfection. The whole-cell protein was obtained by directly lysing the cells in SDS loading buffer. The nucleus protein was obtained by using Nucleoprotein Extraction Kit (Sangon) according to the manufacturer’s instructions. The proteins were denatured by boiling for 10 min, separated by SDS-PAGE, transferred onto a PVDF membrane, and blocked by incubating with Fast Blocking Western buffer (Yeasen). The Cas12f nucleases were detected by incubating with primary mouse-anti-HA mAb (Sangon) and secondary HRP-conjugated goat-anti-mouse mAb (Sangon), and then visualized through chemiluminescence generated by Super ECL Detection Reagent (Yeasen). In parallel, GAPDH and H3C1 were separately detected by incubating with primary mouse-anti-GAPDH (Sangon) and mouse-anti-H3C1 (Sangon) as internal references for whole-cell proteins and nucleus proteins, respectively. To determine the transcript level of sgRNAs cognate to AsCas12f1 and Un1Cas12f1, total RNA was extracted from the transfected cells by using Trizol reagent (Invitrogen). The cDNA was synthesized by using EasyScript All-in-One First-Strand cDNA Synthesis SuperMix (Transgen). Quantitative real-time PCR was then performed using ChamQ SYBR Color qPCR Master Mix (Vazyme) on a LighyCycler 96 system. Relative gene expression was calculated using the 2^−ΔΔCt^ method after normalizing to GAPDH expression.

### Evaluation of genome editing

Genomic DNAs of HEK293T cells were extracted by using Ezup Column Animal Genomic DNA Purification Kit (Sangon). Nearly 200-bp regions surrounding the target loci were amplified using Phanta Max Super-Fidelity DNA Polymerase Mastermix (Vazyme) with primers containing unique sample barcode pairs. PCR products were merged and purified by using TIANquick Midi Purification Kit (TIANGEN). The NGS libraries were prepared by using VAHTS Universal DNA Library Prep Kit for Illumina (Vazyme) according to the manufacturer’s instructions, and then subjected to Illumina Novaseq 6000 sequencing in HaploX Genomics Center (Jiangxi, China). The sequencing data was demultiplexed by using barcodeSpliter (https://github.com/atlasbioinfo/barcodeSpliter) and the indel efficiencies were analyzed by using CRISPResso2 (https://github.com/pinellolab/CRISPResso2)^63^. Information of genomic target sites are listed in Supplementary Dataset 2.

### GUIDE-seq for identifying genome-wide off-target editing

To comprehensively evaluate and compare the genome-wide off-target editing effects among the engineered AsCas12f1-v5.1+T1, Un1Cas12f1-v3.1+ge4.1, enAsCas12a, and SpCas9, GUIDE-seq assays were carried out as previously described^47^. Briefly, the targeting plasmids and dsODN (double-stranded oligodeoxynucleotide) were transfected into HEK293T cells by using Lipofectamine 3000 (Invitrogen) according to the manufacturer’s instructions. The cells were harvested 48-h post transfection for evaluating the genome-wide off-target effects. The genomic DNA was extracted and sheared to an average length of 500 bp by using NEBNext dsDNA Fragmentase (NEB) according to the manufacturer’s instruction. The fragmented gDNA was end-repaired, dA-tailed and ligated to an adapter harboring an 8-nt random molecular index by using VAHTS Universal DNA Library Prep Kit for Illumina V3 (Vazyme) accoding to the manufacturer’s instruction. The dsODN-containing fragments were enriched with two rounds of nested anchored PCR. The library was subjected to Illumina NovaSeq 6000 sequencing in HaploX Genomics Center (Jiangxi, China). Output sequence data was analyzed using the GUIDE pipeline (https://github.com/aryeelab/guideseq).

### Statistics and reproducibility

All statistical analyses were performed using GraphPad Prism (v.9.4.0). The exact replication numbers are indicated in the figure legends. The experimental findings in all of the figures were successfully reproduced at least three times independently.

## Data availability

The structure of AsCas12f1 in complex with sgRNA and target DNA has been deposited in the Protein Data Bank under the accession codes: 7WJU. Source data are provided with this paper.

## Code availability

The sequencing data for evaluating indel efficiencies are demultiplexed by using barcodeSpliter (https://github.com/atlasbioinfo/barcodeSpliter). The indel efficiencies are evaluated by using the CRISPResso2 suite which is available at GitHub (https://github.com/pinellolab/CRISPResso2). The computational pipeline for GUIDE-seq is available at GitHub (https://github.com/aryeelab/guideseq).

## Acknowledgements

This work was supported by grants 2022YFC3400200 from National Key R&D Program of China, LG-QS-202206-05 from the Lingang Laboratory, 22277078, 2207783, and 22207074 from the National Natural Science Foundation of China, 22ZR1480100, and 22YF1428100 from the Shanghai Committee of Science and Technology, China, EKPG21-18 from the Emergency Key Program of Guangzhou Laboratory. The authors also thank the technical support for ICP-OES from the Analytical Instrumentation Center, SPST, ShanghaiTech University.

## Author contribution

Zhaowei Wu, and Q.J. conceived the initial study. Zhaowei Wu, D.L., Z.Z., Y.Q., and H.S. determined the structure of the AsCas12f1 effector complex. Zhaowei Wu, D.P., J.S., J.M., W.F., Zhipeng Wang, F.L., and W.C. performed plasmid construction, protein purification, and biochemical experiments. Zhaowei Wu, D.P., J.S., J.M., W.F. and H.Y. performed the genome editing in human cells and NGS data analyses. Zhaowei Wu, D.P., D.L., X.H., H.S., and Q.J analyzed and discussed the experimental data. Zhaowei Wu and Q.J. prepared the figures and wrote the manuscript. The manuscript was reviewed and approved by all coauthors.

## Competing interests

Q.J., Zhaowei Wu, and D.P. have filed a patent application related to this work through ShanghaiTech University. The remaining authors declare no competing interest.

## Notes

### Competing Interest Statement

Q.J., Zhaowei Wu, and D.P. have filed a patent application (PCT/CN2022/113357) related to this work through ShanghaiTech University. The remaining authors declare no competing interest.

